# Replication protein-A, RPA, plays a pivotal role in the maintenance of recombination checkpoint in yeast meiosis

**DOI:** 10.1101/2023.09.01.555993

**Authors:** Arivarasan Sampathkumar, Zhong Chen, Yuting Tang, Yurika Fujita, Masaru Ito, Akira Shinohara

## Abstract

DNA double-strand breaks (DSBs) activate DNA damage responses (DDR) in both mitotic and meiotic cells. A single-stranded DNA (ssDNA) binding protein, Replication protein-A (RPA) binds to the ssDNA formed at DSBs to activate ATR/Mec1 kinase for the response. Meiotic DSBs induce homologous recombination monitored by a meiotic DDR called the recombination checkpoint that blocks the pachytene exit in meiotic prophase I. In this study, we showed the essential role of RPA in the maintenance of the recombination checkpoint during *S. cerevisiae* meiosis. The depletion of an RPA subunit, Rfa1, in a recombination-defective *dmc1* mutant, fully alleviates the pachytene arrest with the persistent unrepaired DSBs. RPA depletion decreases the activity of a meiosis-specific CHK2 homolog, Mek1 kinase, which in turn activates the Ndt80 transcriptional regulator for pachytene exit. These support the idea that RPA is a sensor of ssDNAs for the activation of meiotic DDR. Rfa1 depletion also accelerates the prophase I delay in the *zip1* mutant defective in both chromosome synapsis and the recombination, suggesting that the accumulation of ssDNAs rather than defective synapsis triggers prophase I delay in the *zip1* mutant.

## Introduction

A eukaryotic single-stranded (ss)DNA binding protein, Replication protein-A (RPA), plays an essential role in DNA replication, DNA repair, and homologous recombination ^1–3^. RPA consists of three subunits; RPA70(RPA1), RPA32(RPA2), and RPA14(RPA3), which are highly conserved from yeast to humans. In *S. cerevisiae*, the knockout of either one of three genes (*RFA1*, *-2* and *-3*) encoding the RPA subunits leads to lethality, consistent with its essential role in DNA replication ^4^. RPA also plays a role in DNA damage response (DDR) ^3,5^.

In homologous recombination. RPA binds to ssDNAs to prevent the formation of the secondary structure of ssDNAs and to protect ssDNAs from degradation by various nucleases ^6^. The assembly of Rad51 filaments on RPA-coated ssDNA is assisted by Rad51 mediators such as Rad52 in the budding yeast ^7–9^. RPA assists Rad51-mediated homology search and strand exchange with a homologous double-stranded (ds)DNA ^10^. The invasion of the ssDNA into the homologous dsDNA results in the formation of a displacement loop (D-loop). The 3’-OH end of the invading strand provides a substrate for DNA synthesis, which is followed by further processing and engaging with the ssDNA in the other DSB end to form a joint molecule. The joint molecule is finally matured into a recombination product. RPA also works in recombination-associated DNA synthesis as well as the annealing in the second-end capture.

DNA double-strand break (DSB) activates the DDR. DSB ends stimulate a DDR kinase, ATM/Tel1 (Ataxia telangiectasia mutated) while ssDNAs do an ATM homolog, ATR/Mec1 (Ataxia telangiectasia mutated related), both of which belong to the PI3 kinase family ^11,12^. The activation of ATR is mediated by the interaction with an ATR partner, ATRIP/Ddc2, which directly interacts with RPA bound to ssDNAs ^13–15^. In mammals, ATR activation is also mediated by the second RPA-binding protein, ETAA1 ^16,17^. Thus, RPA is considered to work as an essential DNA damage sensor to ssDNAs as well as a DNA damage processing protein.

Meiosis rewires DNA transactions such as homologous recombination/DSB repair as well as DDR to guarantee the faithful segregation of two paternal homologous chromosomes at meiosis I for the reduction of chromosome numbers in gametes. In budding yeast, meiotic recombination is initiated by the DSB formation at recombination hotspots by the TopoVI-like protein, Spo11, which is followed by the production of ssDNAs ^18,19^. The ssDNA is used by a meiosis-specific Rad51 homolog, Dmc1 ^20^, for D-loop formation, which is aided by Rad51 ^21^ and its mediator, Mei5-Sae3 complex ^22,23^. Further processing generates a recombination intermediate with double-Holliday junctions (dHJs), which are specifically resolved into crossovers between homologs. Importantly, the partner choice in the recombination is mediated by meiosis-specific chromosome-axis proteins (Hop1, Red1, and Mek1/Mre4), promoting interhomolog bias via an unknown mechanism ^24–26^. Mek1/Mre4 (hereafter Mek1), a meiosis-specific homolog of Rad53/CHK2 kinase ^27,28^, suppresses Rad51-mediated recombination in meiosis ^25,26^. This suppression occurs through the Mek1-mediated phosphorylation of two proteins, Rad54 and Hed1 ^29,30^. Rad54 belongs to the Swi2/Snf2 family protein essential for Rad51-mediated strand invasion ^31^. Mek1-mediated phosphorylation of Rad54 abolishes its interaction with Rad51 ^30^. Hed1, which is expressed only in meiosis, inhibits Rad51 by binding to the Rad51-ssDNA complex ^32^. The inhibitory activity of Hed1 against Rad51 and the stability of Hed1 are augmented by the Mek1-mediated phosphorylation ^29^.

Meiotic events such as recombination and chromosome synapsis are strictly coupled with each other (Fig. 1) ^33^. Similar to mitosis, DSBs in yeast meiosis activate Tel1(ATM) and Mec1(ATR). Different from mitotic DNA damage checkpoint, downstream events of Mec1 or Tel1 activation such as the Rad53 activation are largely suppressed in meiotic cells with poorly-defined mechanisms ^34,35^. On the other hand, Mec1 and/or Tel1 phosphorylates Hop1 on chromosome axis ^36^. Together with chromosomal-bound Red1, phosphorylated Hop1 activates Mek1 kinase, whose autophosphorylation provides positive feedback on its activity ^37^. Mek1-mediated phosphorylation promotes the inactivation of the Ndt80 transcription factor ^38^ essential for the exit from the pachytene stage, where chromosome synapsis and the formation of the dHJs are completed ^37,39^. By activating Ndt80 (through inactivating Mek1), meiotic cells exit the pachytene stage by expressing downstream targets such as Cdc5 Polo-like kinase (PLK) and a cyclin, Clb1 ^40^ for the resolution of dHJs and the disassembly of the synaptonemal complex (SC), a meiosis-specific tripartite chromosome structure, an indicative of chromosome synapsis.

**Figure 1.**
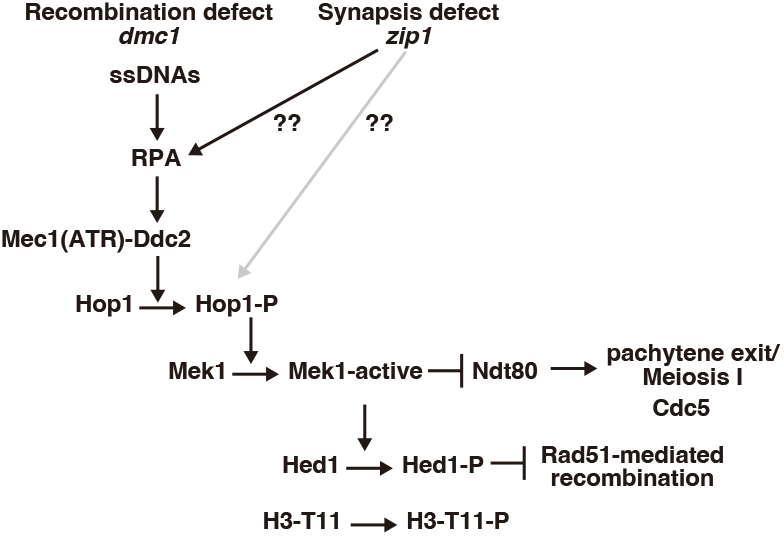
A pathway of recombination checkpoint in meiosis.

It is widely accepted that meiotic cells surveil not only homologous recombination but also chromosome synapsis, referred to as the “recombination” and “synapsis” checkpoints (Fig. 1). Like a recombination-defective *dmc1* mutant, the budding yeast *zip1* mutant, which is defective in the assembly of the central region of the SCs, thus synapsis shows a delay in meiosis progression ^41^. In nematodes and mice, synapsis-defective mutants also delay the meiotic prophase I progression and/or induce apoptosis ^42,43^. In yeast, chromosome synapsis is tightly coupled with meiotic recombination: e.g. a recombination-deficient mutant also displays abnormal synapsis ^44,45^; the *zip1* mutant completely defective in SC formation also impairs the processing of meiotic recombination intermediates. This complicates the presence of the synapsis checkpoint. Moreover, DDR proteins also play a role in meiotic recombination, which also provides a caveat in the interpretation of the meiotic phenotypes of DDR mutants^46–48^.

The role of RPA in DDR is widely accepted. In a pioneer work ^5^, the depletion of RPA70 in human cell lines reduced the ATRIP/Ddc2 binding to the DNA damage as well as the activation of downstream events. In the same study, it was also shown that the depletion of Rfa1 in mitotic yeast cells by a temperature-induced degron reduced the binding of Ddc2 to a single DSB. On the other hand, the other report showed that Rfa1 depletion in G2/M-arrested mitotic cells in budding yeast by the degron does not affect checkpoint activation to a single DSB^6^. A point mutant in the *RFA1* gene encoding RPA70 subunit of yeast RPA called *rfa1-t11* with K45E substitution in the N-terminal protein-protein interaction domain is partially defective in the DDR in the response to telomere dysfunction^49^. Rfa1-t11 showed normal binding to the DSB, but the mutant decreased recruitment of Ddc2 to the DSB ^5^. In meiosis, the *rfa1-t11* with reduced Ddd2-focus formation in nuclei weakly suppresses the *dmc1*-induced delay in meiosis progression ^50^ while RPA containing Rfa1-t11 showed normal localization on meiotic chromosomes ^51^. The *rfa1-t11* mutant alters recombination kinetics ^51^, which may indirectly affect the checkpoint. Furthermore, the conditional depletion of Rpa1 in mouse spermatocytes eliminated RPA foci on meiotic chromosomes but is proficient in ATR recruitment to the chromosomes ^52^. Therefore, we still need careful evaluation of the role of RPA in the DDR in meiotic cells.

In this study, we revisited the role of RPA in the meiotic recombination checkpoint by using a conditional-depletion mutant of Rfa1. The depletion of Rfa1 in the *dmc1*-induced pachytene arrest facilitates the pachytene exit, thus entry into meiosis I and II. Importantly, the RPA depletion did not change the accumulation unrepaired meiotic DSBs in the *dmc1*, which causes abnormal segregation of chromosomes in the division. This confirms the essential role of RPA in the maintenance of the meiotic recombination checkpoint. Moreover, we found that Rfa1 depletion also alleviates delayed meiosis progression in the *zip1* mutant. This suggests that the *zip1* mutation mainly triggers an RPA-dependent response of the prophase I delay, thus the recombination checkpoint rather than the synapsis checkpoint.

## Results

### Depletion of Rfa1 alleviates *dmc1*-induced arrest of meiosis

Three RPA subunits, Rfa1, −2, and −3, are essential for the viability of budding yeast cells ^4^. To know the role of RPA in the DDR in yeast meiosis, we constructed a conditional depletion mutant of the *RFA1* gene encoding the largest RPA subunit, Rfa1, by the auxin-induced degron (AID) system ^53^. The AID tag was added to the C-terminal of the Rfa1 protein. The addition of auxin as well as copper, which induced Tir1 F-box protein, in the medium largely inhibited the mitotic growth of *RFA1-AID* cells (Supplemental Fig. 1a), consistent with the previous study ^4^. *RFA1-AID* diploid cells were induced for the entry of meiosis by incubating in the sporulation medium (SPM). When auxin was added at 4 h incubation with SPM after pre-meiotic DNA replication, which induced the disappearance of Rfa1 from 5 h (Supplemental Fig. 1b), the cells formed few spores after 24 h while more than 90% of the control wild-type cells sporulated (Supplemental Fig. 1c). Most of the Rfa1-depleted cells showed abnormal nuclear morphology (Supplemental Fig. 1c). Indeed, rarely formed tetrad of the cells are all dead (0 viable spores in 100 tetrads) while without the auxin treatment, the viability of spores from *RFA1-AID* cells was 91.8% (among 100 tetrads). The pulse-field gel electrophoresis revealed the accumulation of meiotic DSBs on chromosomes (Supplemental Fig. 1d), confirming a defect in DSB repair ^51^. These show that the AID-tag did not interfere with Rfa1 function in both miotic and meiotic cells and that Rfa1 is essential for both mitosis and meiosis.

To know the role of RPA in the recombination checkpoint in meiosis, we used the *dmc1* mutant, which is unable to repair meiotic DSBs and, as a result, shows an arrest at mid-pachytene stage during meiotic prophase I ^20^. This response to unrepaired DSBs is referred to as the recombination checkpoint. *RFA1-AID* was introduced in the *dmc1* deletion mutant; *dmc1 RFA1-AID*. Like the *dmc1* mutant, without auxin addition, *dmc1 RFA1-AID* shows an arrest in meiotic prophase I without division (e.g. Fig. 2d), indicating that the Rfa1 tagging did not affect the recombination checkpoint in the absence of Dmc1. RPA is also involved in DNA replication and S-phase checkpoint ^1,2^, which indirectly affects the G2 event. To avoid the effect in pre-meiotic DNA replication, we depleted Rfa1 protein by adding auxin to a meiotic culture of *dmc1 RFA1-AID* cells at 4 h after the induction of meiosis in the sporulation medium (SPM), thus, after pre-meiotic DNA replication, which occurs at 1.5-3 h. At 5 h, after the 1-hour addition of auxin, the protein level of Rfa1-AID was decreased to ∼15% level relative to the original level at 4 h (Fig. 2a, Supplementary Figs. S1e and S2), indicating the efficient depletion of Rfa1 under the condition. We also checked the focus formation of Rfa2, the second subunit of RPA, on meiotic chromosome spreads. Rfa2 shows punctate staining on the spreads with other recombination proteins such as Rad51 ^54^, which corresponds with ongoing recombination (Fig. 2b). In the *dmc1* mutant, Rfa2 focus-staining appears at 2.5 h, reaches a plateau at 5 h, and persists after that, indicating little turnover of Rfa2 due to stalled recombination in the mutant (Fig. 2c, Supplementary Fig. S3a). When auxin was added at 4 h, Rfa2 foci disappeared within one hour, and Rfa2 foci were rarely observed during further incubation by 10 h (Fig. 2b, c). This suggests that RPA is highly dynamic and easily dissociated from the chromosome. We concluded that Rfa1 depletion at 4 h in meiosis eliminates most chromosome-bound RPA complexes by 5 h.

**Figure 2.**
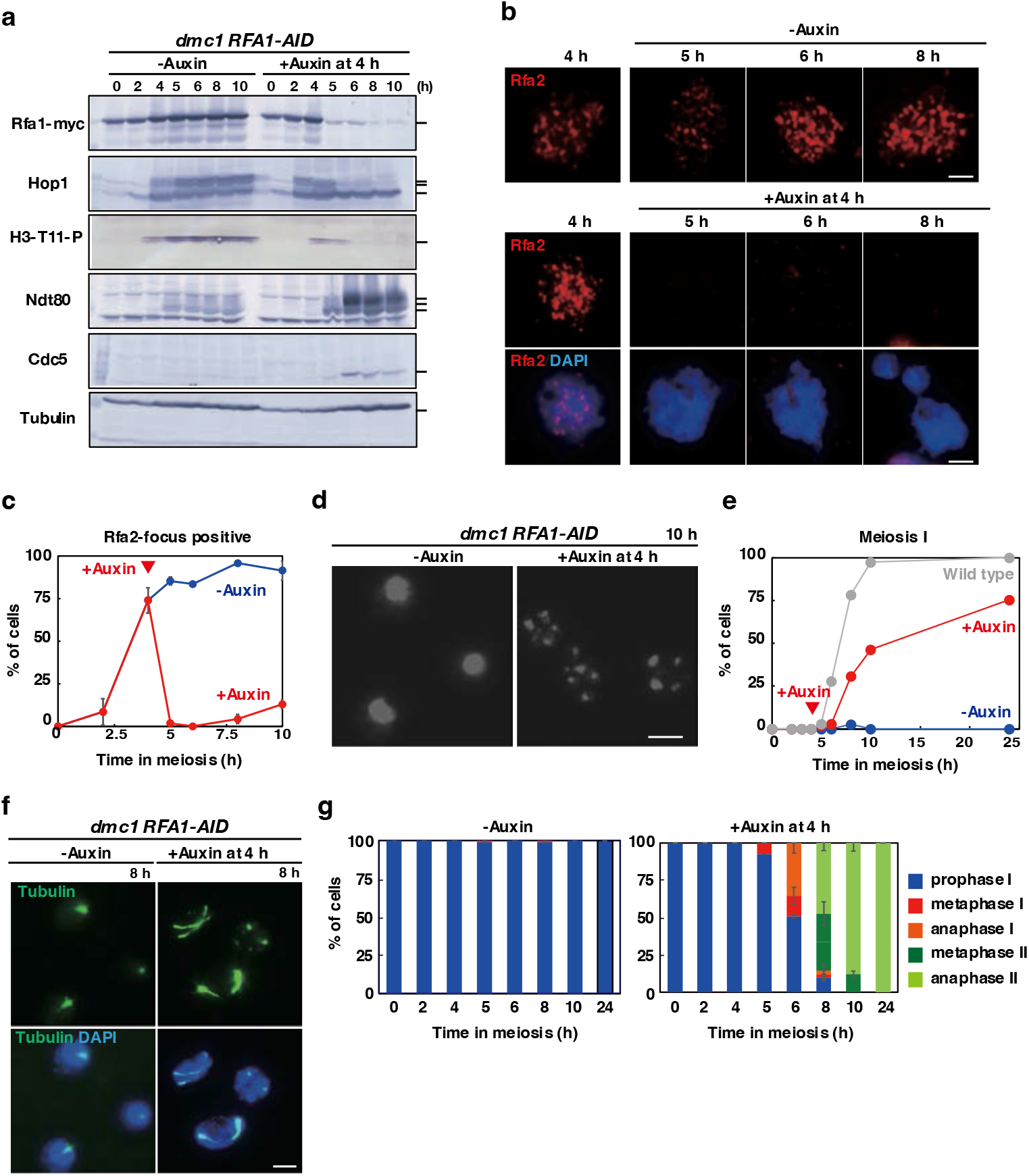
Rfa1 depletion alleviates *dmc1*-induced pachytene arrest. (a) Expression of various proteins in meiosis. Lysates obtained from the *dmc1 RFA1-AID* (SAY63/64) cells at various time points during meiosis with the addition of auxin (2 mM) in DMSO or DMSO alone at 4 h were analyzed by western blotting using anti-myc (Rfa1-myc-AID, upper), anti-Hop1, anti-histone H3-phospho-T11, anti-Ndt80, anti-Cdc5 or anti-tubulin (lower) antibodies. (b) Rfa2 staining. Nuclear spreads from *dmc1 RFA1-AID* (SAY63/64) cells with or without the addition of auxin at 4 h were stained with anti-Rfa2 (red), and DAPI (blue). Representative images at each time point under the two conditions are shown. Bar = 2 μm. (c) Kinetics of Rfa2 assembly/disassembly. The number of Rfa2-positive cells (with more than 5 foci) was counted at each time point. At each time point, more than 100 cells were counted. (d) Meiosis progression. *dmc1 RFA1-AID* (SAY63/64) cells were analyzed by DAPI staining. Representative image of cells treated with/without 4 h-auxin addition at 10 h are shown. Bar = 2 μm. (e) Meiosis progression. The entry into meiosis I and II in *dmc1 RFA1-AID* (SAY63/64) cells were analyzed by DAPI staining. The number of DAPI bodies in a cell was counted. A cell with 2, 3, and 4, and 3 and 4 DAPI bodies was defined as a cell that passed through meiosis I and meiosis II, respectively. The graph shows the percentages of cells that completed MI or MII at the indicated time points. More than 200 cells were counted at each time point. The representative results (*n*=3) are shown. *dmc1 RFA1-AID* without auxin (with DMSO), blue closed circles; *dmc1 RFA1-AID* with auxin addition at 4 h, red closed circles; Wild type (MSY823/833), gray circles. (f) Whole cells were fixed and stained with anti-tubulin (green) and DAPI (blue). Representative images at 8 h of *dmc1 RFA1-AID* (SAY63/64) cells with or without the addition of auxin at 4 h are shown. Bar = 2 μm. (g) Classification of tubulin staining at each time point of meiosis in *dmc1 RFA1-AID* (SAY63/64) cells with (right) or without (left) the addition of auxin at 4 h. Dot, short line, and long line tubulin-staining with single DAPI mass were defined as prophase I (blue), metaphase I (red), anaphase I (orange), metaphase II (green) and anaphase II (light green). At each time point, more than 100 cells were counted. The representative results are shown (*n*=3).

### Checkpoint alleviation in Rfa1 depletion in *dmc1* meiosis

DAPI staining of DNAs confirmed a single DAPI body in a *dmc1 RFA1-AID* cell even at 24 h, thus an arrest in prophase I as shown previously ^20^. Rfa1 depletion at 4 h induced fragmentation of DAPI bodies at late time points. In some cells, 2-4 irregular-sized DAPI bodies with smaller DAPI bodies were detected in a single cell (Fig. 2d). The cells with abnormal DAPI bodies appeared at 6 h and gradually accumulated by 24 h (Fig. 2e). This suggests the presence of nuclear division, thus the onset of meiosis I and/or II in the *dmc1* cells upon Rfa1 depletion. We also checked the effect of Rfa1 depletion on meiotic DNA replication. The FACS analysis showed that the addition of auxin at 4 h did not affect the transition from 2C to 4C (Supplementary Fig. S1f).

To evaluate the meiosis progression precisely, we examined the status of spindle morphology in the *dmc1 RFA1-AID* cells in the absence or presence of auxin (Fig. 2f, g). Tubulin staining revealed a single focus associated with single or multiple short projections in meiotic prophase I of the cell in the absence of auxin (Fig. 2f). The *dmc1 RFA1-AID* mutant maintained this arrangement of spindles in the absence of auxin by 24 h, consistent with an arrest of the *dmc1* mutant in prophase I (Fig. 2f). On the other hand, the Rfa1 depletion induced the elongation of spindles after one hour of addition (at 5 h). At 6 h, ∼50% *dmc1 RFA1-AID* cells contained single short and long spindles corresponding with metaphase I and anaphase I, respectively (Fig. 2e). At 8 h, in addition to the metaphase I and anaphase I spindle, ∼70% of cells contained metaphase II or anaphase II spindles (Fig. 2f, g). At 10 h, most of the cells showed diffuse staining of microtubules in multiple DAPI bodies in a cell. These show that Rfa1 depletion almost alleviates meiotic prophase I arrest induced by the *dmc1* mutation.

Cdc5/PLK is expressed after the pachytene exit ^40,55^. Like *dmc1* cells, without auxin addition, the *dmc1 RFA1-AID* mutant cells did not express Cdc5 (Fig. 2a). On the other hand, the addition of auxin at 4 h induced the expression of Cdc5 from 5 h. This confirms the exit of the pachytene stage of *dmc1* cells in the absence of RPA.

### No DSB repair in *dmc1* meiosis with Rfa1 depletion

One possibility to suppress the *dmc1* arrest is to channel unrepaired DSBs into the repair pathway. On the other hand, the acute depletion of RPA did not affect the status of meiotic DSBs under *dmc1* arrest. The CHEF (Counter-clamped homogeneous electric field) gel electrophoresis confirmed the accumulation of fragmented chromosomes in the *dmc1* mutant irrespective of the presence or absence of RPA. In the *dmc1 RFA1-AID* mutant without auxin treatment, smear bands of full-length chromosomes appeared at 3 h and most of the chromosomes became fragmented with a size of less than 700 kbp by 5 h. After 6 h, fragmented chromosomes persisted by 24 h (Fig. 3a, Supplementary Fig. S4 for wild-type control). The depletion of Rfa1 from 4 h did not affect the persistence of fragmented chromosomes. This suggests that meiotic DSBs remain unrepaired under the depletion of RPA in the *dmc1* mutant.

**Figure 3.**
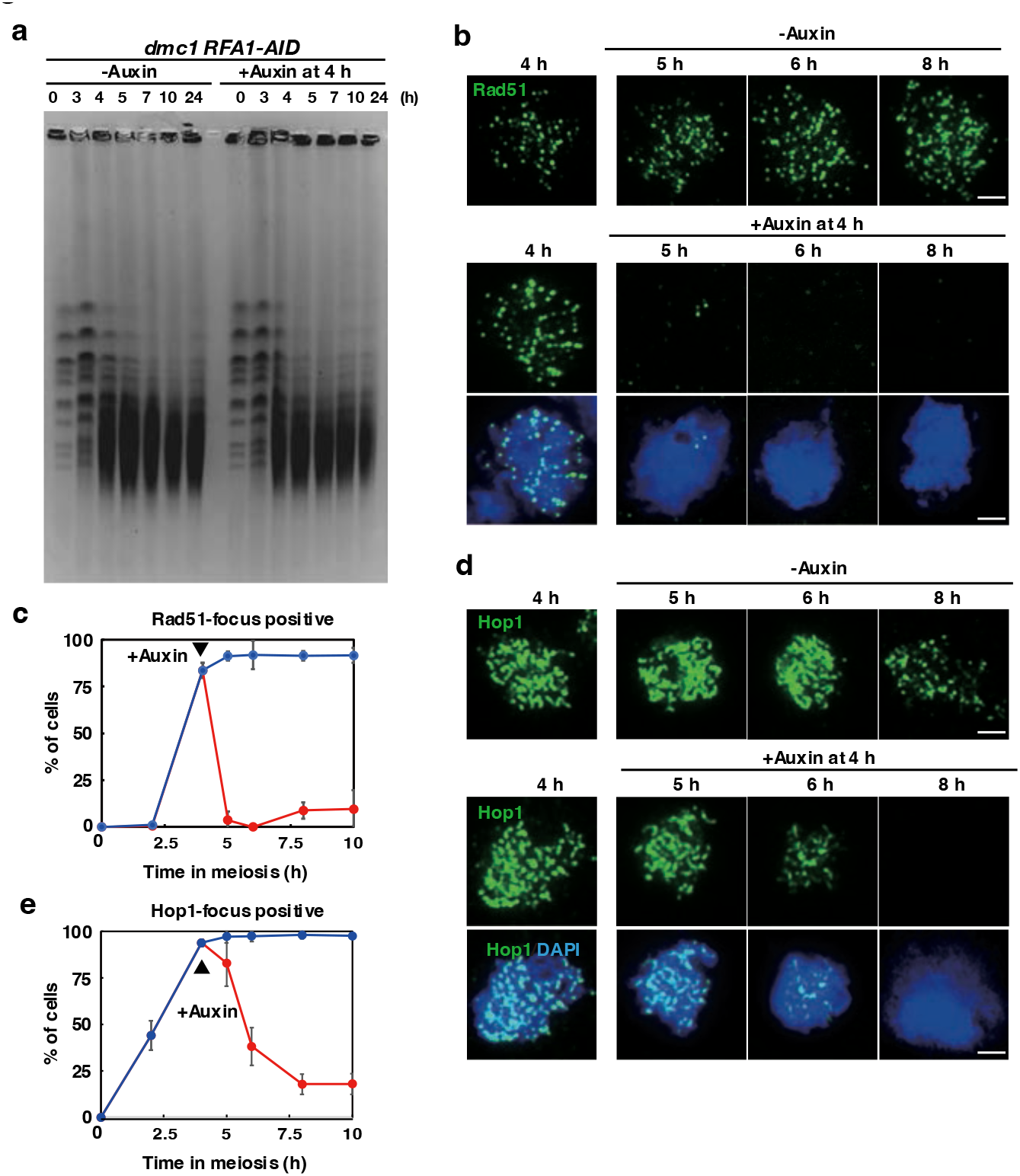
Rfa1 depletion does not affect DSB turnover in *dmc1* mutant. (a) CHEF analysis of meiotic DSB repair. Chromosomal DNAs from *dmc1 RFA1-AID* (SAY63/64) cells with the addition of auxin (2 mM) in DMSO or DMSO alone at 4 h were studied by CHEF electrophoresis. The gel is stained with EtBr. Auxin was added at 4 h in the incubation with SPM. (b) Rad51 staining. Nuclear spreads from *dmc1 RFA1-AID* (SAY63/64) cells with or without the addition of auxin at 4 h were stained with anti-Rad51 (green), and DAPI (blue). Representative images at each time point under the two conditions are shown. Bar = 2 μm. (c) Kinetics of Rad51 assembly/disassembly. The number of Rad51-positive cells (with more than 5 foci) was counted at each time point. At each time point, more than 100 cells were counted. (d) Hop1 staining. Nuclear spreads from *dmc1 RFA1-AID* (SAY63/64) cells with or without the addition of auxin at 4 h were stained with anti-Hop1 (green), and DAPI (blue). Representative images at each time point under the two conditions are shown. Bar = 2 μm. (e) Kinetics of Hop1 assembly/disassembly. The number of Rad51-positive cells (with more than 5 foci) was counted at each time point. At each time point, more than 100 cells were counted.

We also indirectly checked the DSB status by analyzing Rad51 foci on meiotic chromosomes (Fig. 3b, c). Like Rfa2 foci, in the *dmc1 RFA1-AID* mutant in the absence of auxin, Rad51 foci appeared at 4 h, reached a plateau at 5 h, and persisted at late times as shown previously ^56^. The addition of auxin at 4 h rapidly induced the disappearance of Rad51 foci at 5 h. The reduced Rad51 foci are seen at late time points (Fig. 3c and Supplementary Fig. 3b). This suggests that RPA is critical for the maintenance of Rad51 foci on the chromosomes. The absence of Rad51 foci in the Rfa1 depletion supports the inability of *dmc1 RFA1-AID* cells without RPA to repair meiotic DSBs by Rad51-dependent recombination.

### The Hop1/Mek1-Ndt80 axis was down-regulated after RFA1 depletion

Un-repaired DSBs in the *dmc1* mutant activate the recombination checkpoint, which down-regulates a meiosis-specific transcription activator, Ndt80 for the pachytene exit (Fig. 1). DSB-activated Mec1/ATR kinase phosphorylates a HORMAD protein, Hop1, which stimulates a meiosis-specific kinase, Mek1. Mek1 binds and phosphorylates Ndt80 to inactivate it. For pachytene exit, further phosphorylation of Nd80 by Ime2 kinase promotes the activation of Ndt80 ^38^. In *dmc1 RFA1-AID* cells, the Ndt80 appeared at 4 h with multiple bands and accumulated more during further incubation in SPM without a change of band-shift status (Fig. 2a). At 6 h, after two hours of addition of auxin, a slower-migrating Ndt80 band appeared and accumulated. Importantly, as shown above, under the depletion condition, Cdc5/PLK, which is a major target of activated Ndt80 and is necessary for pachytene exit, appeared at 6 h (Fig. 2a). This is consistent with meiosis progression of the *dmc1* mutant cells induced by Rfa1-depletion.

We also checked the behavior of the upstream regulators of Ndt80, and Hop1 by western blotting. In the *dmc1* mutant, Hop1 appeared as a non-phosphorylated form at 2 h and phosphorylated Hop1 with slower migrated bands persisted from 4 h (Fig. 2a) as shown previously ^36^. After Rfa1 depletion, major bands of phosphorylated Hop1 almost disappeared from 6 h with a similar steady-state level of non-phosphorylated Hop1 during the time course. This suggests that Hop1 is de-phosphorylated during pachytene exit. On chromosome spreads, Hop1 shows beads-in-line staining for the localization of chromosome axes (Fig. 3d) ^57,58^. In the *dmc1* mutant without Rfa1 depletion, Hop1 localization remains unchanged during meiosis (Fig. 3d). The addition of auxin at 4 h induced the disappearance of Hop1-positive spreads from 5 h. At 6 h, ∼35% of cells are positive for Hop1 while ∼95% are positive in a control (Fig. 3d, e). At 8 h, most of the Hop1 signals on the chromosomes disappeared.

One of the Mek1 kinase targets is histone H3T11 phosphorylation ^59^. Under the *dmc1* arrest condition, H3T11 phosphorylation accumulated (Fig. 2a). On the other hand, the Rfa1 depletion rapidly induced the disappearance of the signal (Fig. 2a), suggesting the downregulation of Mek1 activity under Rfa1 depletion. These results clearly showed that Rfa1, thus RPA, is critical for the maintenance of the recombination checkpoint through the Hop1/Mek1-Ndt80 axis.

The *dmc1*-induced arrest in meiosis is also partially suppressed by the *rad51* ^60^. A recent study showed that the suppression of *dmc1*-arrest by the *rad51* mutation is mediated by partial elongation of SCs with concomitant loss of Hop1^61^. Rad51 loss from the chromosomes in the *dmc1* mutant upon Rfa1-depletion such as 5-h time point by 4-h addition of auxin seems to mimic a condition seen in the *rad51 dmc1* double mutant. To address whether Rad51-focus loss in the *dmc1* mutant by Rfa1 depletion induced SC formation, we checked SC formation by staining Zip1, a component of SC central region ^41^. The Zip1 staining is classified into three classes; dots, short lines, and long lines. As shown previously ^20^, the *dmc1* mutant is defective in SC elongation (Supplemental Fig. 5a, b). Rfa1 depletion promotes the SC disassembly by the pachytene exit. At 5 h when most of Rad51 foci disappeared, we did not see elongation of Zip1 lines relative to control with Rad51 foci and rather observed increased Zip1 polycomplex, which is indicative of defective SC assembly (Supplemental Fig. 5c). Moreover, Hop1 localization one-hour after Rfa1 depletion is similar to that without the depletion (Fig. 3d, e). These suggests that the suppression of arrest by Rfa1 depletion is unlikely to come from altered Hop1 functions as seen in the *rad51 dmc1* double mutant ^61^.

### Bypass of *dmc1* arrest by Rfa1 depletion depends on Ndt80

We tested whether the pachytene exit of the *dmc1* cells by Rfa1 depletion depends on the *NDT80* (Fig. 4). The addition of auxin to *dmc1 ndt80 RFA1-AID* cells efficiently reduced the amount of Rfa1 protein (Fig. 4a, Supplementary Fig. S6) and Rfa2 foci from meiotic chromosomes (Fig. 4b). However, different from the *dmc1 RFA1-AID* mutant, the addition of auxin to *dmc1 ndt80 RFA1-AID* mutant did not induce the elongation of spindles (Fig. 4c) or Hop1 disassembly from the chromosomes (Fig. 4d). Thus, the pachytene exit of *dmc1* cells by Rfa1 depletion still depends on Ndt80. Interestingly, the amount of the phosphorylated Hop1 is reduced slightly compared to that in the control (Fig. 4a). This is less drastic compared to the presence of Ndt80, supporting an auto-feedback mechanism between Mek1 activation and Ndt80 activation ^62^.

**Figure 4.**
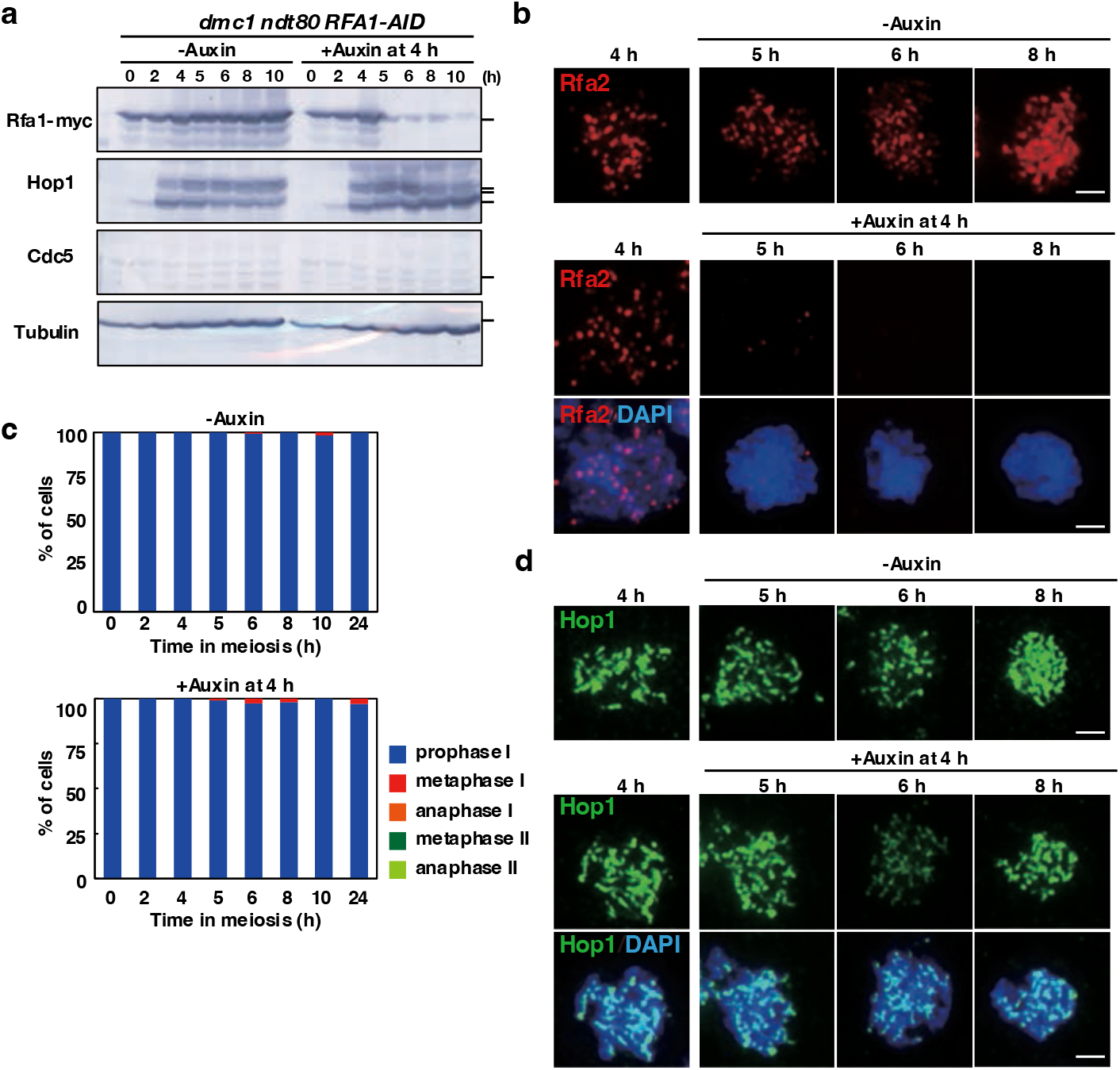
*NDT80* is necessary for pachytene exit by Rfa1 depletion. (a) Expression of various protein in meiosis. Lysates obtained from the *ndt80 dmc1 RFA1-AID* (SAY80/81) cells at various time points during meiosis with the addition of auxin (2 mM) in DMSO or DMSO alone were analyzed by western blotting using anti-myc (Rfa1-myc-AID, upper), anti-Hop1, anti-Cdc5 or anti-tubulin (lower) antibodies. (b) Rfa2 staining. Nuclear spreads from *ndt80 dmc1 RFA1-AID* (SAY80/81) cells with or without the addition of auxin at 4 h were stained with anti-Rfa2 (red), and DAPI (blue). Representative images at each time point under the two conditions are shown. Bar = 2 μm. (c) Classification of tubulin staining at each time point of meiosis in *ndt80 dmc1 RFA1-AID* (SAY80/81) cells with (right) or without (left) the addition of auxin at 4 h. Dot, short line, and long line tubulin-staining with single DAPI mass were defined as prophase I (blue), metaphase I (red), anaphase I (orange), metaphase II (blue), anaphase II (green) and post-meiosis II (pale green). Short and long tubulin-staining were defined as metaphase II and anaphase II, respectively. At each time point, more than 100 cells were counted. The representative results are shown (*n*=2). (d) Hop1 staining. Nuclear spreads from *ndt80 dmc1 RFA1-AID* (SAY SAY80/81) cells with or without the addition of auxin at 4 h were stained with anti-Hop1 (green), and DAPI (blue). Representative images at each time point under the two conditions are shown. Bar = 2 μm.

### Checkpoint alleviation in Rfa1 depletion in *zip1* meiosis

Meiotic yeast cells monitor not only ongoing meiotic recombination but also chromosome synapsis ^43^. However, the presence of the so-called “synapsis checkpoint” in yeast meiosis is under debate ^62^. The mutant in the *ZIP1* gene is deficient in chromosome synapsis with a delay of the entry into meiosis I and is used to analyze the synaptic checkpoint ^41^. We examined the effect of Rfa1 depletion on the *zip1*-induced delay of meiosis I (Fig. 5). As reported previously ^45^, in the SK1 strain background, *zip1 RFA1-AID* cells without auxin addition started meiosis I at 8 h while wild-type cells entered at 5 h (and finished meiosis II by 10 h) (Fig. 5b). When auxin was added at 4 h, which induced rapid depletion of Rfa1 and the dissociation of Rfa2 from meiotic chromosomes (Fig. 5a, c, h, Supplementary Fig. S7), metaphase I spindles appeared at 5 h, and at 6 h, ∼35% of the cells contained metaphase-I and anaphase-I spindles (Fig. 5d, e). At 8 h, ∼30% of the cells contained anaphase-II spindle. Importantly, DAPI staining showed the entry of the *zip1* cells into meiosis I upon Rfa1 depletion is similar to that in wild-type cells (Fig. 5b). The spore viability of *zip1 RFA1-AID* cells treated with auxin was 0% while cells without auxin was 35.5% (among 100 tetrads).

**Figure 5.**
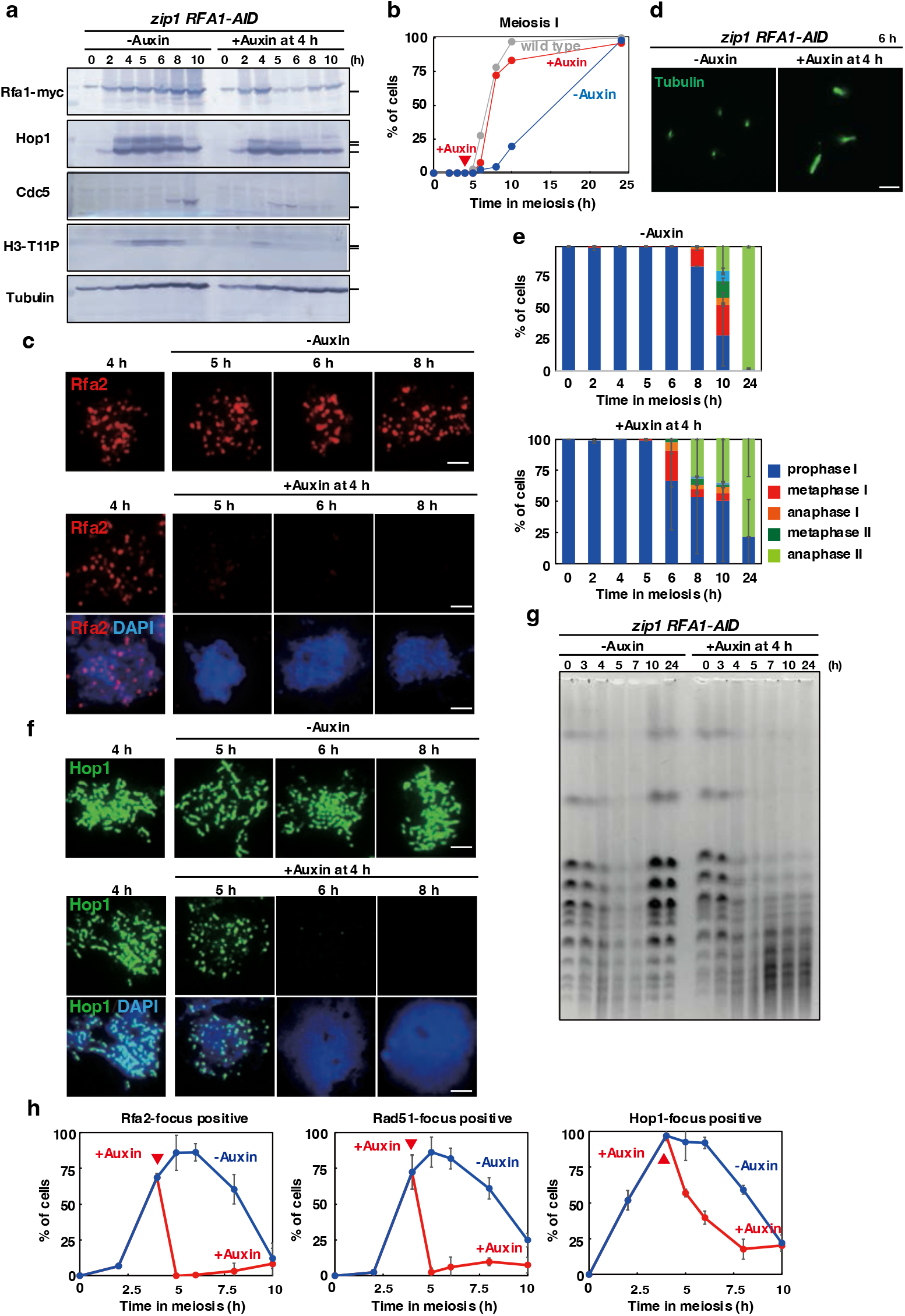
Rfa1 depletion alleviates *zip1*-induced delay in meiosis I. (a) Expression of various proteins in meiosis. Lysates obtained from the *zip1 RFA1-AID* (SAY72/73) cells at various time points during meiosis with the addition of auxin (2 mM) in DMSO or DMSO alone were analyzed by western blotting using anti-myc (Rfa1-myc-AID, upper), anti-Hop1, anti-Cdc5, anti-histone H3-phospho-T11 or anti-tubulin (lower) antibodies. (b) Meiotic cell cycle progression. The entry into meiosis I and II in *zip1 RFA1-AID* (SAY72/73) cells were analyzed by DAPI staining as shown in Figure 2D. More than 200 cells were counted at each time point. The representative results (*n*=3) are shown. *dmc1 RFA1-AID* without auxin, blue closed circles; *dmc1 RFA1-AID* with auxin addition at 4 h, red closed circles. (c) Rfa2 staining. Nuclear spreads from *zip1 RFA1-AID* (SAY72/73) cells with or without the addition of auxin at 4 h were stained with anti-Rfa2 (red), and DAPI (blue). Representative images at each time point under the two conditions are shown. Bar = 2 μm. (d) Tubulin staining in the *zip1 RFA1-AID* mutant. Whole cells were fixed and stained with anti-tubulin (green) and DAPI (blue). Representative images for each class are shown. Bar = 2 μm. (e) Classification of tubulin staining at each time point of meiosis in *zip1 RFA1-AID* (SAY72/73) cells with (right) or without (left) the addition of auxin at 4 h. The quantification was performed as shown in Figure 1G. At each time point, more than 100 cells were counted. The representative results are shown (*n*=2). (f) Hop1 staining. Nuclear spreads from *zip1 RFA1-AID* (SAY72/73) cells with or without the addition of auxin at 4 h were stained with anti-Hop1 (green), and DAPI (blue). Representative images at each time point under the two conditions are shown. Bar = 2 μm. (g) CHEF analysis of meiotic DSB repair. Chromosomal DNAs from *zip1 RFA1-AID* (SAY72/73) cells with or without the addition of auxin at 4 h were studied by CHEF electrophoresis. Auxin was added at 4 h in the incubation with SPM. (h) Kinetics of assembly/disassembly of Rfa2 (left), Rad51 (middle), and Hop1 (right). The number of focus-positive cells (with more than 5 foci) was counted for Rfa2 and Rad51 at each time point. At each time point, more than 100 cells were counted.

The *zip1 RFA1-AID* cells without auxin expressed Cdc5 from 8 h (Fig. 5a), consistent with delayed progression of prophase I in the mutant. The addition of auxin at 4 h accelerated the Cdc5 expression from 5 h. The *zip1* mutant accumulated H3T11 phosphorylation of histone H3 signal from 4 h to 8 h (Fig. 5a). The Rfa1 depletion accelerated the disappearance of this phosphorylation from 5 h. The accelerated disappearance of the phospho-Hop1 protein is also seen by the depletion (Fig. 5a, f). These showed the Rfa1-dependent inactivation of the Hop1/Mek1-Ndt80 signaling pathway induced by the *zip1* mutation. Thus, the delay induced by the *zip1* mutation largely depends on the Rfa1.

In addition to SC formation, Zip1 plays a critical role in meiotic recombination as a pro-crossover factor. Indeed, the *zip1* mutant shows delayed DSB repair. The CHEF analysis showed the presence of broken chromosomes in the *zip1* mutant from 3 h to 7 h (Fig. 5g). Importantly, full-length chromosomes were reassembled at 10 h without Rfa1 depletion, indicating that most of the DSBs in the *zip1* mutant were repaired with the delay. On the other hand, Rfa1 depletion from 4 h induced fragmentation of the chromosomes from 5 h. Even after 24 h, little recovery of full-length chromosomes was seen under the condition. This suggests that Rfa1 is essential for meiotic DSB repair in the *zip1* mutant.

### Rfa1 depletion does not alleviate delay induced by *rad50S* meiosis

The *rad50S* mutant, which is defective in the end-processing of meiotic DSBs, also shows delayed meiosis I progression ^63^. This delay in the *rad50S* mutant is mediated by Tel1(ATM) and Rad50-Mre11-Xrs2, which recognize DSB ends ^64^. The pathway is distinct from the Mec1(ATR)-mediated pathway shown above. We also examined the effect of Rfa1 depletion on the *rad50S*-induced delay (Fig. 6a, Supplementary Fig. S8). As reported, without auxin, *rad50S RFA1-AID* cells delay the onset of meiosis I as shown by whole-cell staining of tubulin (Fig. 6b). Rather, the addition of auxin to the *rad50S RFA1-AID* cells delayed the onset of meiosis I more than without the auxin. We cannot detect Rfa2 foci in the r*ad50S* mutant even in the presence of Rfa1 (Fig. 6c), confirming that the *rad50S* accumulates unresected DSBs ^63^. Moreover, Hop1 phosphorylation is not observed in the mutant (Fig. 6a). This is consistent with little formation of ssDNAs without DSB processing in the mutant ^63^, thus little Mec1/ATR activation. This is expected since the *rad50S* mutant does not form any ssDNAs, which is a substrate for RPA. The accumulation of chromosomal Hop1 in the mutant is not affected by the Rfa1 depletion (Fig. 6d).

**Figure 6.**
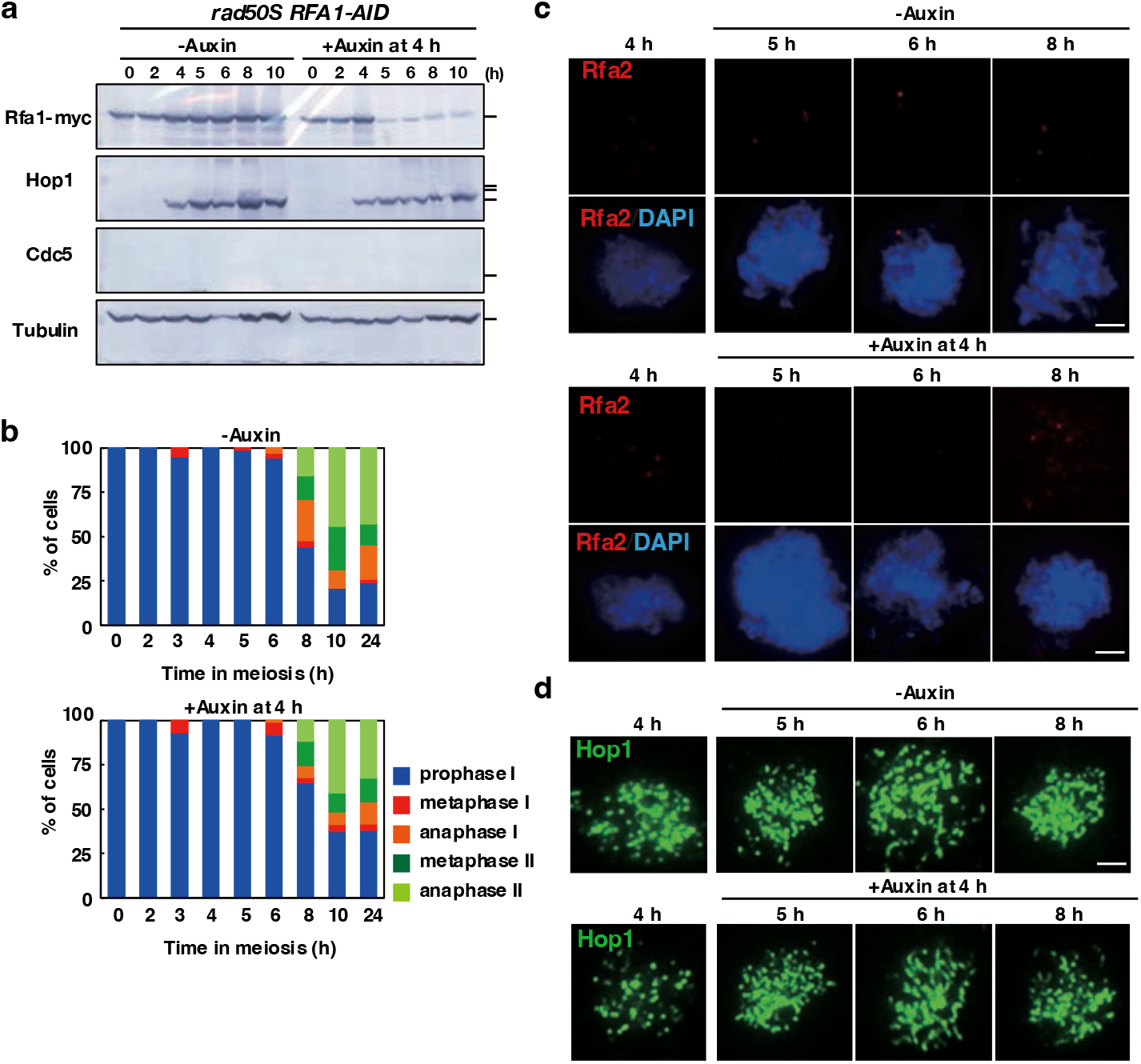
Rfa1 depletion does not suppress *rad50S*-induced delay in meiosis I. (a) Expression of various proteins in meiosis. Lysates obtained from the *rad50S RFA1-AID* (SAY68/69) cells at various time points during meiosis with the addition of auxin (2 mM) in DMSO or DMSO alone were analyzed by western blotting using anti-myc (Rfa1-myc-AID, upper), anti-Hop1, anti-Cdc5 or anti-tubulin (lower) antibodies. (b) Classification of tubulin staining at each time point of meiosis in *rad50S RFA1-AID* (SAY68/69) cells with (right) or without (left) the addition of auxin at 4 h. The quantification was performed as shown in Figure 1G. At each time point, more than 100 cells were counted. The representative results are shown (*n*=1). (c) Rfa2 staining. Nuclear spreads from *rad50S RFA1-AID* (SAY68/69) cells with or without the addition of auxin at 4 h were stained with anti-Rfa2 (red), and DAPI (blue). Representative images at each time point under the two conditions are shown. Bar = 2 μm. (d) Hop1 staining. Nuclear spreads from *rad50S RFA1-AID* (SAY68/69) cells with or without the addition of auxin at 4 h were stained with anti-Hop1 (red), and DAPI (blue). Representative images at each time point under the two conditions are shown. Bar = 2 μm.

## Discussion

RPA plays multiple roles in DNA replication, DNA repair, and recombination through its ability to bind to ssDNAs with a high affinity. RPA is also important for DDR as a sensor of DNA damage, thus ssDNAs. The role of RPA in Mec1(ATR) activation through the binding of Mec1(ATR)-interaction protein, Ddc2(ATRIP), to RPA has been documented well ^5,13,14^. In this paper, we showed the pivotal role of RPA in the maintenance of DDR in meiosis, called recombination checkpoint.

### The role of RPA in recombination checkpoint

Previously, in the budding yeast, two *RFA1*-degron mutants, *rfa1-td* and *td-RFA1*, were constructed and characterized to study the role of Rfa1 in DDR activation. Indeed, the *rfa1-td*, as well as the *rft1-t11 (rfa1-K45E)* mutant, reduce the recruitments of Ddc2 to a DSB site ^5^. On the other hand, it is shown that, by using the *td-RFA1*, G2/M-arrested haploid cells depleted for Rfa1 still activate Rad53 upon a single DSB ^6^. The latter study concludes that a low level of RPA is sufficient for the activation of Mec1-Ddc2. More recently, an RPA-independent pathway for ATR activation, which requires APE1, was identified in *Xenopus* egg extracts ^65^. We revisited the role of RPA in DDR by studying the effect of Rfa1 depletion in DDR such as the meiotic recombination checkpoint. The recombination checkpoint in meiotic cells responds to the ssDNA through RPA-Mec1-Ddc2 axis as seen in a canonical DNA damage checkpoint in mitotic cells. By using the auxin-degron, we depleted Rfa1 in the *dmc1* mutant which arrests in meiotic prophase I with hyper-resected ssDNAs in ∼150 DSBs ^20^. The Rfa1-depletion induced the pachytene exit by downregulating Mec1 and Mek1 activities, which in turn activates the Ndt80-dependent gene expression of key regulators for the pachytene exit such as Cdc5 ^40,55^. This is consistent with the previous study which showed that the *rfa1-t11* with reduced Ddd2-focus formation in nuclei weakly suppresses the *dmc1*-induced delay ^50^. Importantly, the Rfa1 depletion did not affect the accumulation of ssDNAs in the *dmc1* cells with persistent unrepaired DSBs. This is particularly important since mutations in DDR genes such as *RAD17*, *-24*, *DDC1*, and *MEC3* even in the *MEC1* gene bypass the *dmc1*-induced arrest by repairing DSBs by an alternative pathway such as ectopic recombination ^46,48,66^. The results described here supported the idea that RPA is a sensor for ssDNAs, which creates a scaffold for the assembly of checkpoint effectors such as Mec1-Ddc2(ATR-ATRIP). In meiosis, meiosis-specific Mek1 is activated through the phosphorylation of Hop1 by Mec1 kinase^36^. The Rfa1-depletion induces rapid dephosphorylation of Hop1 and the dissociation from meiotic chromosomes without affecting the protein level, suggesting the presence of an active de-phosphorylation pathway of Hop1.

### The chromosome synapsis checkpoint in yeast meiosis

In addition to the recombination checkpoint, meiotic cells responds to abnormal chromosome synapsis, which is called synapsis checkpoint. This idea was proposed that mutants defective in chromosome synapsis show a delay in the progression of prophase I and apoptosis in yeasts and mice or nematodes, respectively. However, the presence of the synapsis checkpoint is still under debate ^42,43^, since in most organisms meiotic recombination and chromosome synapsis are tightly coupled. The budding yeast *zip1* mutant, which is completely defective in SC formation, thus, chromosome synapsis, delays or arrest in meiotic prophase I ^41,45^. This arrest is alleviated by the mutant of genes encoding a component of checkpoint such as *PCH2* ^67^. Pch2 modulates Hop1 conformation and regulates the association and dissociation of Hop1 on meiotic chromosomes ^61,68,69^. On the other hand, the *zip1* mutant also shows defective DSB repair and/or the crossing-over ^44,45^, which suggests delayed progression of meiosis I in the *zip1* mutant is caused by the impaired recombination. In this study, by using the *RFA1-AID*, we could test whether the delay in *zip1* mutant depends on RPA or not and found that the Rfa1-depletion eliminates the delay induced by the *zip1* mutation. This strongly suggests that the recombination checkpoint delays the meiosis I progression in the *zip1* mutant as seen in the *dmc1* mutant. Moreover, consistent with this, the other yeast mutants, which are fully defective in the synapsis with a weak recombination defect such as *ecm11* and *gmc2*, do not show a drastic delay in the meiosis I progression ^70,71^. Thus, at least in budding yeast which couples the recombination and chromosome synapsis, rather than the status of chromosome synapsis, a recombination process, the presence of ssDNAs, can be monitored by a meiosis-specific surveillance mechanism to guarantee the completion of meiotic recombination directly and chromosome synapsis indirectly. Our idea is consistent with the idea proposed by recent reports claiming the merging of the recombination and synapsis checkpoints in meiosis ^61,72^.

## Methods

### Strains and plasmids

All strains described here are derivatives of *S. cerevisiae* SK1 diploids, MSY832/833 (*MATα/MAT**a**, lys2/’’, ura3/’’, leu2::hisG/’’, trp1::hisG/’’*). The genotypes of each strain used in this study are described in Supplementary Table S1

### Strain Construction

The *RFA1-AID* gene was constructed by one-step DNA integration. The AID-9Myc tag was added to the C-terminus of the *RFA1* open reading frame by the transformation of yeast cells (MSY823) with a DNA fragment amplified by PCR using p7AID-9myc (provided by Dr. Neil Hunter) as a template and a pair of primers (Rfa1-AID-Hygoro-F and -R in Supplementary Table S2). The transformants were selected against hygromycin. Genomic DNAs from the candidate colonies were purified and the genomic DNAs were checked for the correct tagging by PCR and DNA sequencing using the primer described in Supplementary Table S2.

### Anti-serum and antibodies

Anti-myc (9E10, sc-40, Santa Cruz), anti-tubulin (MCA77G, Bio-Rad/Serotec, Ltd), anti-Cdc5 (sc-33625, Santa Cruz), and histone H3 phospho-T11 (ab5168, Abcam) were used for western blotting. Rabbit anti-Rad51 ^21^ and guinea pig anti-Rad51 ^73^, rabbit anti-Hop1 ^74^ and rabbit anti-Rfa2 ^75^ were home-made and used for staining. Rabbit anti-Ndt80 was a gift by M. Lichten. The secondary antibodies for immuno-staining were Alexa Fluor 488 (Goat, A-21206, Thermo Fisher Scientific) and 594 (Goat, A-211012, Thermo Fisher Scientific) IgG used at a 1/2000 dilution. For western blotting, alkaline phosphatase-conjugated secondary antibodies (Promega) were used.

### Meiotic Time course

*Saccharomyces cerevisiae* SK1 strains were patched onto YPG plates (2% bacteriological peptone, 1% yeast extract, 3% glycerol, 2% bacteriological agar) and incubated at 30 °C for 12 h. Cells were inoculated onto YPD plates (2% bacteriological peptone, 1% yeast extract, 2% glucose, 2% bacteriological agar) and grown for two days to obtain isolated colonies. A single colony was inoculated into 3 ml of YPD liquid medium and grown to saturation at 30 °C overnight. To synchronize cell cultures, the overnight cultures were transferred to pre-warmed SPS medium (1% potassium acetate, 1% bacteriological peptone, 0.5% yeast extract, 0.17% yeast nitrogen base with ammonium sulfate and without amino acids, 0.5% ammonium sulfate, 0.05 M potassium biphthalate) and grown for 16– 17 h. Meiosis was induced by transferring the SPS-cultured cells to pre-warmed SPM (1% potassium acetate, 0.02% raffinose). The SPM-cultured cells were harvested at various times after transfer. For Rfa1 depletion from 4 h, CuSO_4_ was added to 50 mM at 3.5 h. Auxin (3-indoleacetic acid) solved in dimethyl sulfoxide (DMSO) was added at a final concentration of 2 mM at 4 h. In the control experiment (without auxin), we added an equivalent DMSO solution alone.

### Cytology

Chromosome spreads were immunostained as described previously ^48,73^. Spheroplasts were burst in the presence of 1% paraformaldehyde and 0.1% lipsol. The spreads were stained with primary and secondary antibodies observed using a fluorescence microscope (BX51; Olympus/Evident) with a 100× objective (NA1.3). Images were captured by CCD camera (CoolSNAP; Roper), and then processed using IP lab and/or iVision (Bio Vision Technologies) and Photoshop (version, 25.4.0; Adobe; https://www.adobe.com) software. For focus counting, ∼30 nuclei were counted at each time point.

Whole-cell staining was carried out as follows. Yeast cells are fixed with formaldehyde (a final concentration of 3.7%) and collected. The pellets were suspended in the ZK buffer (25 mM Tris-HCl [pH 7.5], 80 mM KCl) and treated with DTT (20 mM) and then with Zymolyase 100T (2.5 μg/mL). After washing with PBS, the cells were placed on a glass slide coated with poly-lysine (1 mg/ml). The cells were treated with cold ethanol (100% 6 min) and acetone (100% 0.5 min), and washed with PBS. After blocking with PBS with 1% BSA, cells were stained with primary and secondary antibodies.

### Western blotting

Western blotting was performed as described previously ^22,76^, using cell lysates extracted by the TCA method. After being harvested and washed twice with 20% TCA, cells were roughly disrupted with zirconia beads by the Multi-beads shocker (Yasui Kikai). The protein precipitate recovered by centrifugation was suspended in SDS-PAGE sample buffer adjusted to pH 8.8 and then incubated at 95 °C for 10 min. After electrophoresis, the proteins were transferred onto a nylon membrane (Immobilon-P, Millipore) and incubated with primary antibodies in a blocking buffer (1× PBS, 0.5% BSA) and then with alkaline phosphatase-conjugated secondary antibody (Promega). The color reaction was developed with NBT/BCIP solution (Nacalai Tesque).

### CHEF analysis Pulsed-field gel electrophoresis

For pulsed-field gel electrophoresis (PFGE), chromosomal DNA was prepared in 1.3% of low-melting point agarose plugs as described in ^74^ and run at 14 °C with 1X TBE buffer in a CHEF DR-III apparatus (BioRad) using the field 6V/cm at a 120°angle for 48 h. Switching times followed a ramp from 15.1 to 25.1 seconds. After the electrophoresis, the gel was stained with 0.5 μg/mL ethidium bromide for 30 min and washed with distilled water. Images were taken using ImageQuant LAS4000 BioImager (GE Healthcare).

### FACS analysis

Cells in 1 ml of SPM culture were collected by centrifugation. The pellet was then resuspended in 1 ml of 70% ethanol. Fixed cells were recollected and washed by suspending 1 ml of 50 mM sodium citrate buffer. The collected cells were resuspended in 0.5 ml of sodium citrate buffer and sonicated for the 20s at room temperature. Sonicated cells were incubated with 12.5 μl of 10 mg/mL RNAseA for 1 h at 50 °C and in 25 μl of 15mg/ml proteinase K for 1h at 50 °C. 0.5ml of 16 μlg/mL PI solution was added and incubated for 30 min at room temperature in the dark. The samples were then analyzed using SA-3800 Spectral analyzer (Sony). Floreada.io was used for gating and plotting graphs.

## Data availability

Uncropped western blotting images are provided in Supplementary Figures. The raw data used in plots and graphs are also shown in Source Data. The raw microscopy images used for figures and quantification are available from corresponding authors upon request.

## Acknowledgments

We are grateful to Drs. Neil Hunter for providing the yeast AID system and Michael Lichten for anti-Ndt80. A.Sa. is a JICA fellow. Y.F. and M.I. were supported by the Institute for Protein Research. This work was supported by a grant from Institute for Fermentation, Osaka (IFO) and from Uehara Memorial Foundation to Life Sciences; to A.Sh.

## Author contributions

A.Sh. conceived and designed the experiments. A.Sa., Z.C., and Y.T. carried out experiments with help from Y.F. and M.I.. A.Sa, Z.C., Y.T., Y.F., M.I., and A.Sh. analyzed the data. A.Sh. wrote the manuscript with inputs from help from others.

