## Supplemental Information for "Replication protein-A, RPA, plays a pivotal role in the maintenance of recombination checkpoint in yeast meiosis"

Supplementary Table 1-2
Supplementary Figure 1-8

**Table S1. Strain List**

| Strain number | Genotypes |
|---|---|
| MSY831/833 | *MAT**a**/* $\alpha$, *ho::LYS2/", lys2/", ura3/", leu2::hisG/", trp1::hisG/"* |
| NKY1551 | *MAT**a**/* $\alpha$, *ho::LYS2/", lys2/", ura3/", leu2::hisG/", his4X-LEU2(BamHI)-URA3/his4B-LEU2(MluI), arg4-bgl/arg4-nsp* |
| SAY15/16 | MSY831/833 with *Rfa1-AID::HYG/", p-CUP1-Os-Tir1::URA3* |
| SAY63/64 | SAY15/16 with *dmc1::URA3/"* |
| SAY68/69 | SAY15/16 with *rad50S::LEU2/"* |
| SAY72/73 | SAY15/16 with *zip1::LEU2/"* |
| SAY80/81 | SAY15/16 with *ndt80::LEU2/"* |

**Table S2. Oligonucleotide list**

| Primer name | Sequence |
|---|---|
| Rfa1-AID-Hygro-F | 5’-TTG AAT TAC AGG GCT GAA GCC GAC TAT CTT GCC GAT GAG TTA TCC AAG GCT TTG TTA GCT CGT ACG CTG CAG GTC GAC |
| Rfa1-AID-Hygro-R | 5’-TTT TTT TTT TAC ATT TCT CAT ATG TTA CAT AGA TTA AAT AGT ACT TGA TTA TTT GAT ACA ATC GAT GAA TTC GAG CTC G |
| RFA1-F | 5’-GTG AGA AGT GCG ACA CCA ATA |
| RFA1-R | 5’-GTC GAC TAT TTG GAG AAG GAA G |

### Supplementary Figure S1. Arivarasan S. et al.

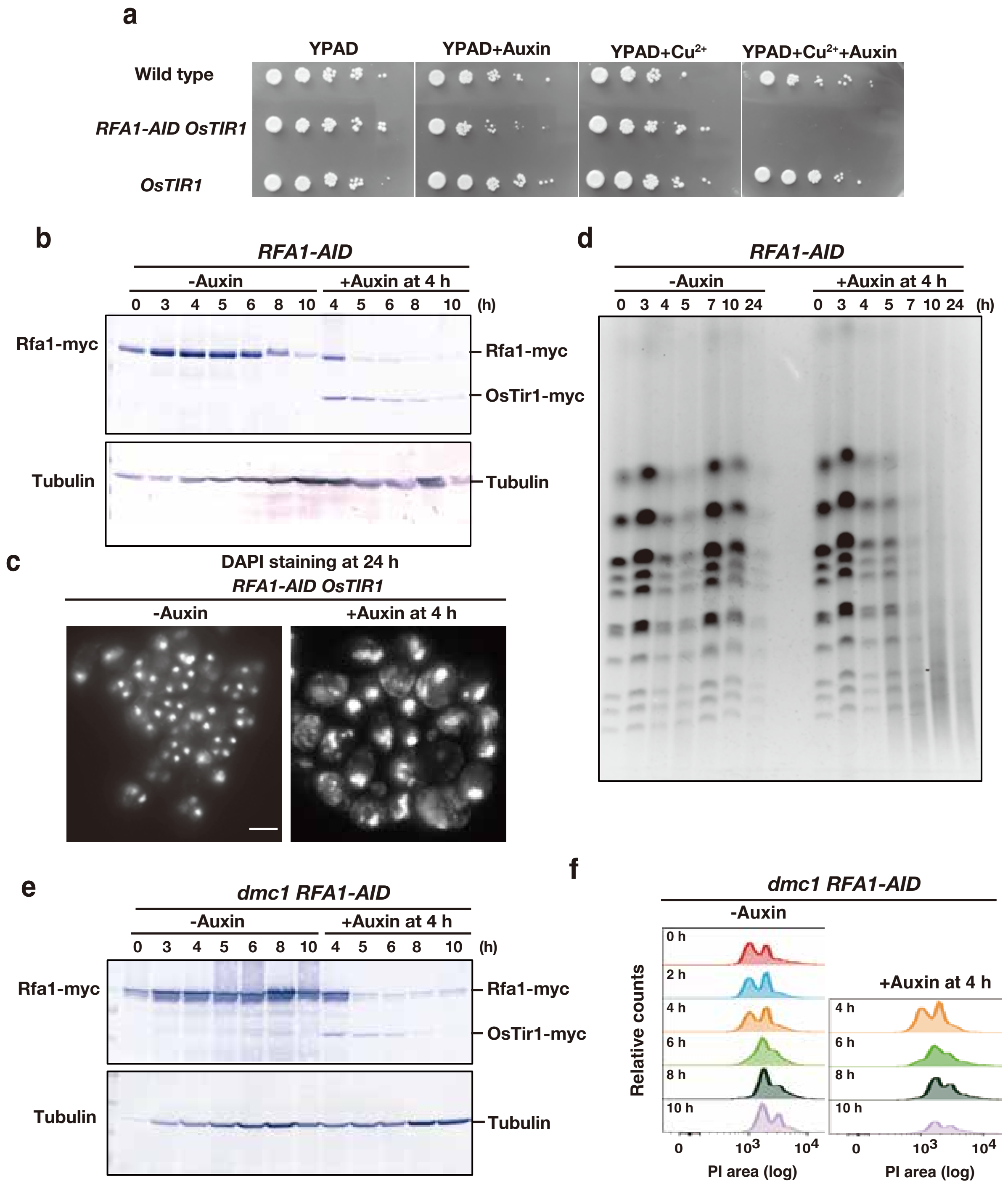

**Supplementary Figure S1.**

(a) Viability of *RFA1-AID* diploid (SAY15/16) cells. 10-fold serially diluted diploid cells were spotted on YPAD plates with or without auxin and $CuSO_4$ and incubated at 30 °C for 48 h.

(b) Expression of Rfa1-AID protein in meiosis. Lysates obtained from the *RFA1-AID* (15/16) cells at various time points during meiosis with the addition of auxin (2 mM) in DMSO or DMSO alone at 4 h were analyzed by western blotting using anti-myc (Rfa1-myc-AID and OsTir1-myc, upper), or anti-tubulin (lower) antibodies.

(c) DAPI-stained image of *RFA1-AID* diploid cells (SAY15/16) after incubating in SPM for 24 h. Auxin was added at 4 h in the incubation of SPM. Representative images are shown. Bar = 2 μm.

(d) CHEF analysis of meiotic DSB repair. Chromosomal DNAs from *dmc1 RFA1-AID* (SAY15/16) cells with or without the addition of auxin at 4 h were studied by CHEF electrophoresis. Auxin was added at 4 h in the incubation with SPM.

(e) Expression of Rfa1-AID protein in meiosis. Lysates obtained from the *dmc1 RFA1-AID* (SAY63/64) cells at various time points during meiosis with or without the addition of auxin (2 mM) at 4 h were analyzed by western blotting using anti-myc (Rfa1-myc-AID and OsTir1-myc, upper), or anti-tubulin (lower) antibodies.

(f) FACS profile of the *dmc1 RFA1-AID* mutant (SAY63/64) without or with auxin addition at 4 h. Fixed cells at each time point were stained with PI and analyzed by FACS cell sorter.

**Supplementary Figure S2. Arivarasan S. et al.**

#### Figure 2a uncropped blot

##### Anti-myc (for Rfa1-myc)

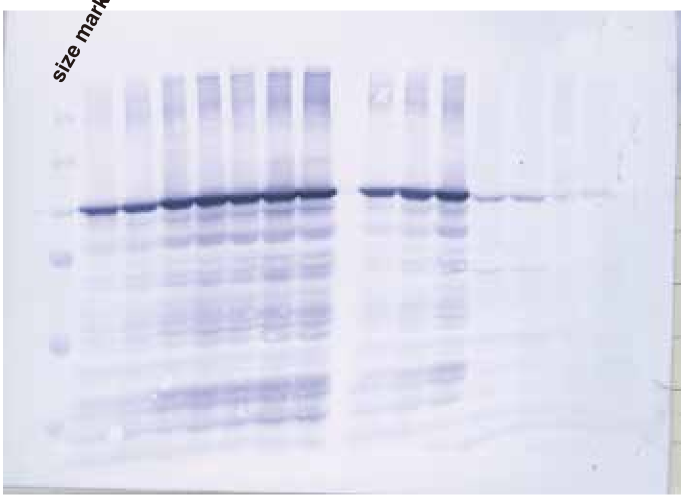


##### Anti-Hop1

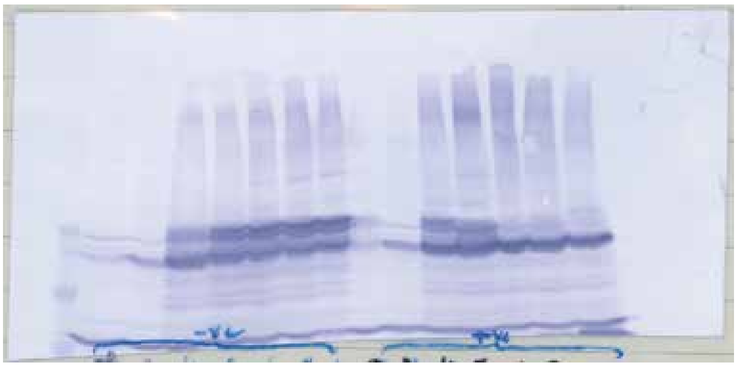

##### Anti-Histone H3-phosphoT11

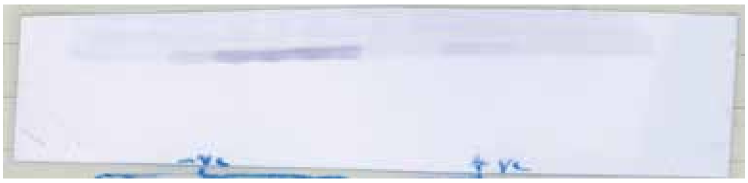

##### Anti-Cdc5

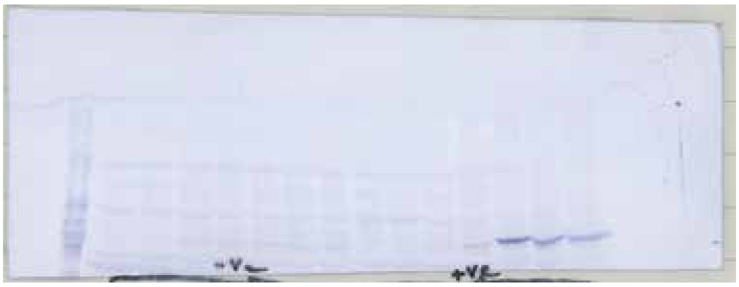

##### Anti-Ndt80

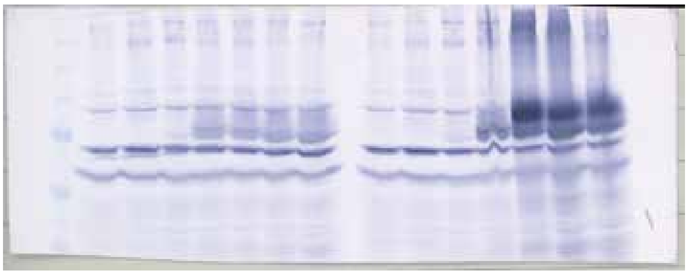

##### Anti-Tubulin

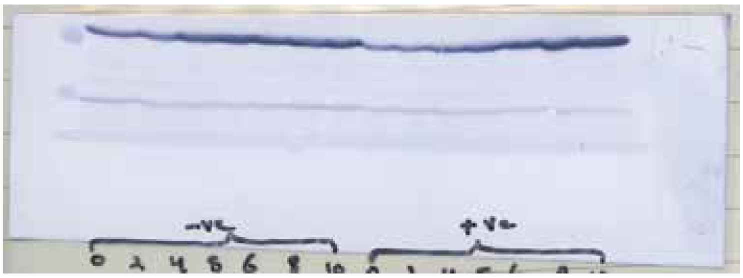

**Supplementary Figure S2.**

Uncropped images of western blotting in Figure 2a.

#### Supplementary Figure S3. Arivarasan S. et al.

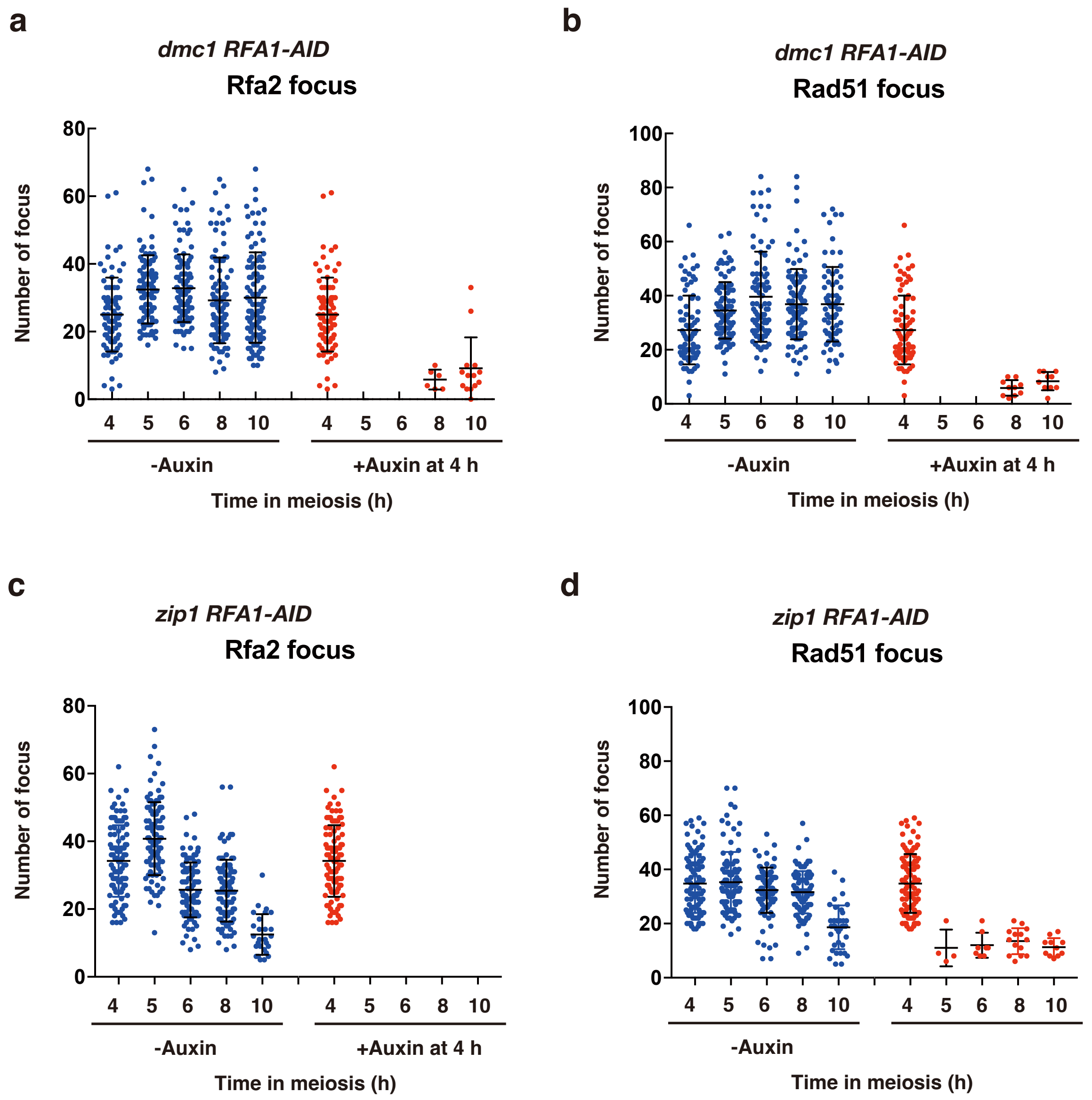

**Supplementary Figure S3.**

(a) The number of Rfa2 foci per spread in *dmc1 RFA1-AID* (SAY72/73) cells with (red) or without (blue) the addition of auxin at 4 h was counted at different time points. The mean and SD are shown in the plot.

(b) The number of Rad51 foci per spread *dmc1 RFA1-AID* (SAY72/73) cells with (red) or without (blue) the addition of auxin at 4 h was counted at different time points. The mean and SD are shown in the plot.

(c) The number of Rfa2 foci per spread in *zip1 RFA1-AID* (SAY72/73) cells with (red) or without (blue) the addition of auxin at 4 h was counted at different time points. The mean and SD are shown in the plot.

(d) The number of Rad51 foci per spread *zip1 RFA1-AID* (SAY72/73) cells with (red) or without (blue) the addition of auxin at 4 h was counted at different time points. The mean and SD are shown in the plot.

**Supplementary Figure S4. Arivarasan S. et al.**

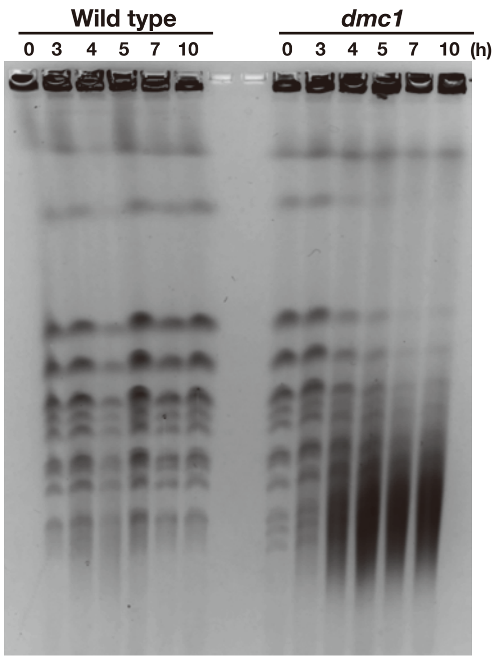

**Supplementary Figure S4.**

CHEF analysis of meiotic DSB repair. Chromosomal DNAs from Wild type (MSY831/833) and the *dmc1* mutant cells were studied by CHEF electrophoresis. Auxin was added at 4 h in the incubation with SPM.

#### Supplementary Figure S5. Arivarasan S. et al.

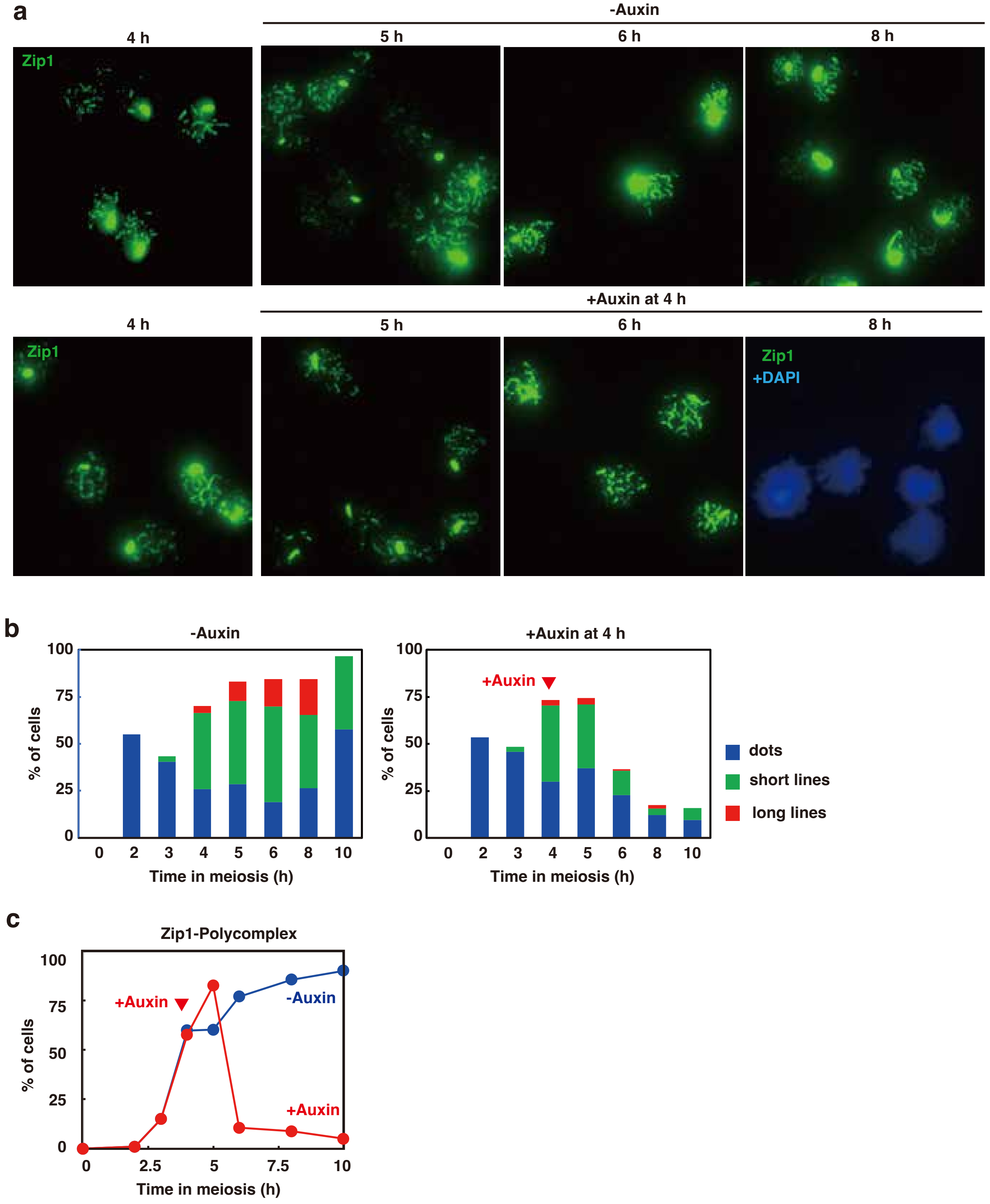

**Supplementary Figure S5.**

(a) Zip1 staining. Nuclear spreads from *dmc1 RFA1-AID* (SAY63/64) cells with or without the addition of auxin at 4 h were stained with anti-Zip1 (green), and DAPI (blue). Representative images at each time point under the two conditions are shown. Bar = 2 μm.

(b) Classification of Zip1 staining at each time point of meiosis in *dmc1 RFA1-AID* (SAY63/64) cells with (right) or without (left) the addition of auxin at 4 h. Dot, short line, and long line tubulin-staining with single DAPI mass were defined as prophase I (blue), short lines (green), and long lines (red). At each time point, more than 100 cells were counted. The averages are shown (*n*=2).

(c) Kinetics of Zip1 polycomplex. The number of cells with Zip1 polycomplex was counted at each time point. At each time point, more than 100 cells were counted. *dmc1 RFA1-AID* (SAY63/64) cells with (red) or without (blue) the addition of auxin at 4 h.

**Supplementary Figure S6. Arivarasan S. et al.**

#### Figure 4a uncropped blot

**Anti-myc (for Rfa1-myc)**

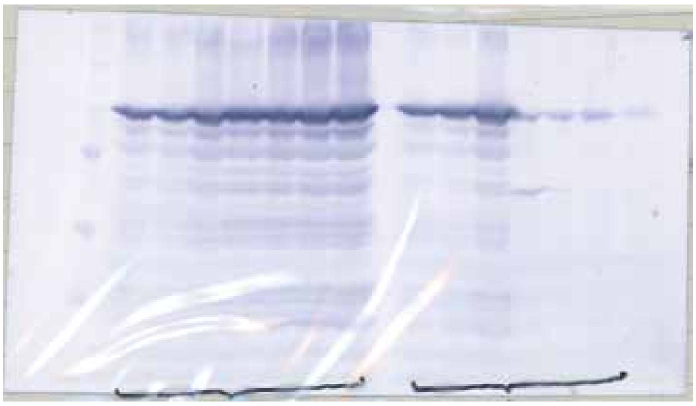

**Anti-Hop1**

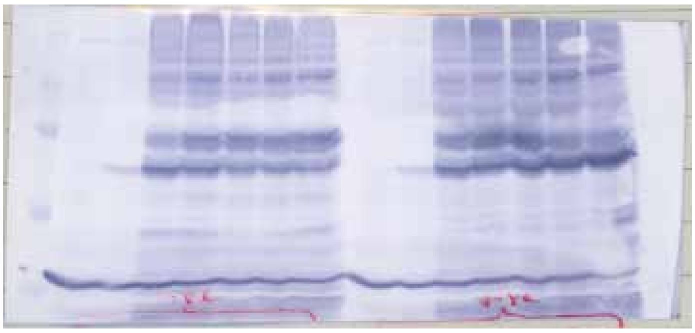

**Anti-Cdc5**

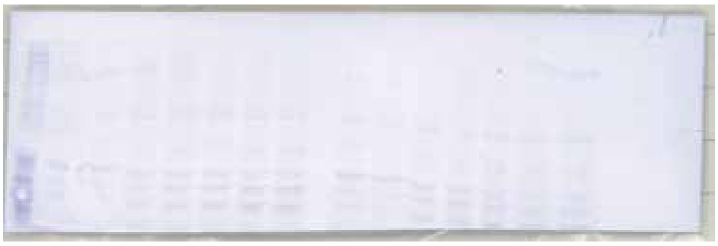

**Anti-Tubulin**

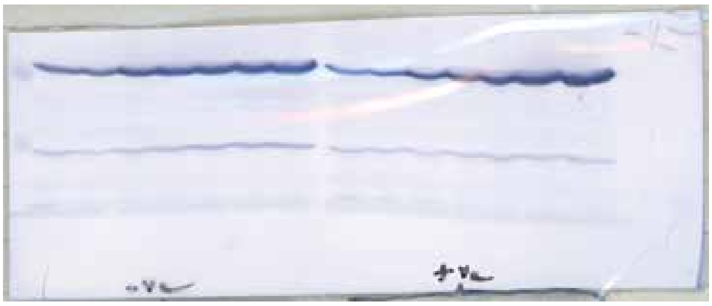

**Supplementary Figure S6.**

Uncropped images of western blotting in Figure 4a.

Supplementary Figure S7. Arivarasan S. et al.

#### Figure 5a uncropped blot

##### Anti-myc (for Rfa1-myc)

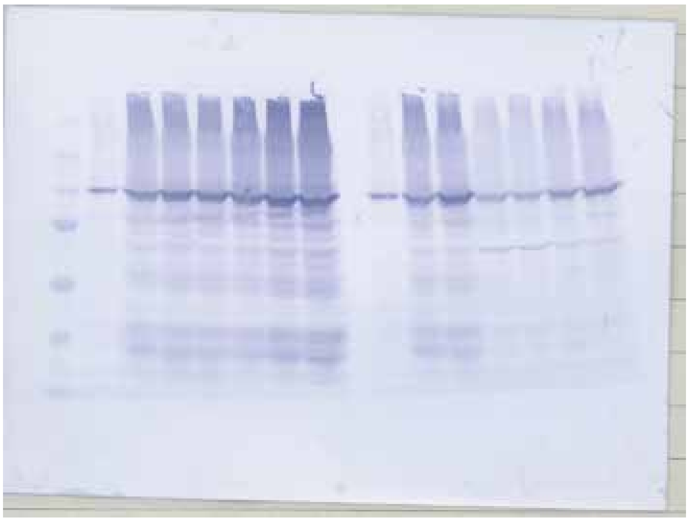

##### Anti-Cdc5

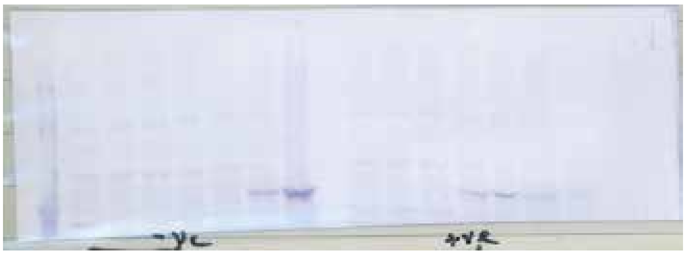


##### Anti-Tubulin

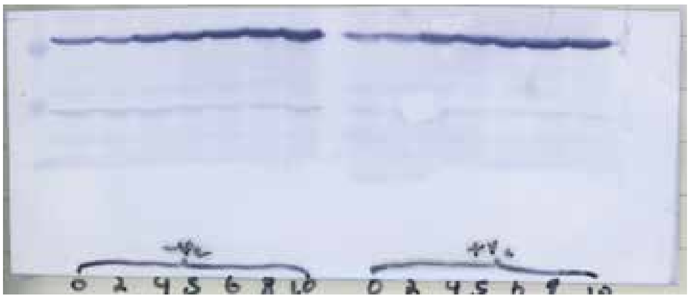


##### Anti-Hop1

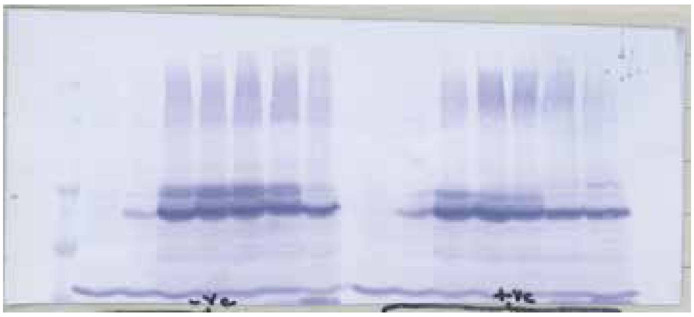


##### Anti-Histone H3-phosphoT11

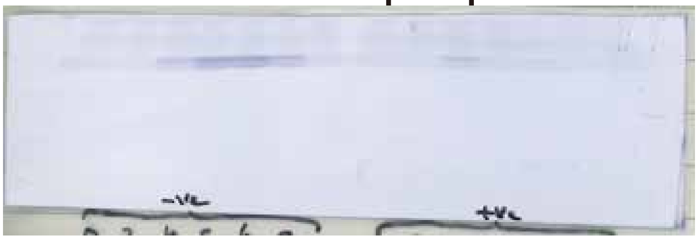

**Supplementary Figure S7.**

Uncropped images of western blotting in Figure 5a.

**Supplementary Figure S8. Arivarasan S. et al.**

#### Figure 6a uncropped blot

##### Anti-myc (for Rfa1-myc)

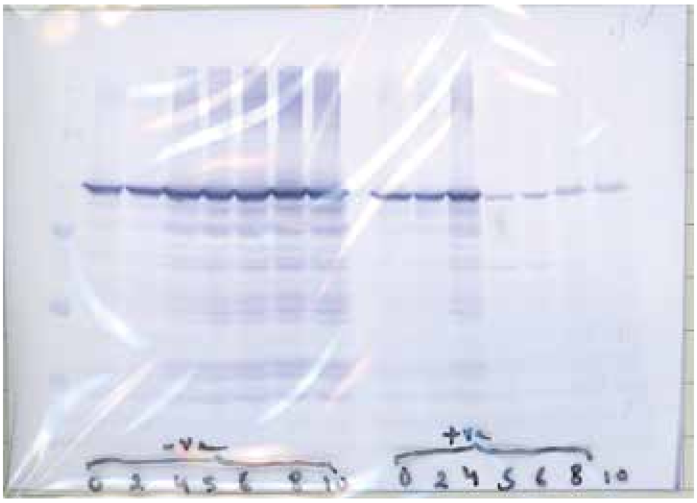


##### Anti-Hop1

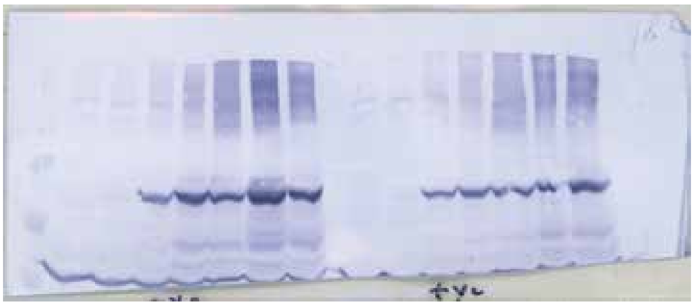


##### Anti-Cdc5

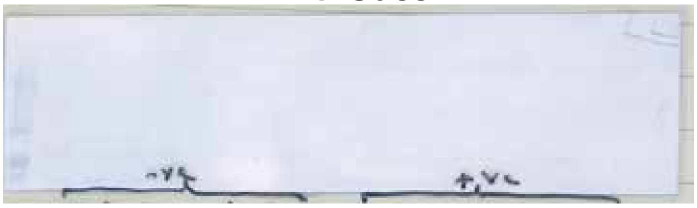


##### Anti-Tubulin

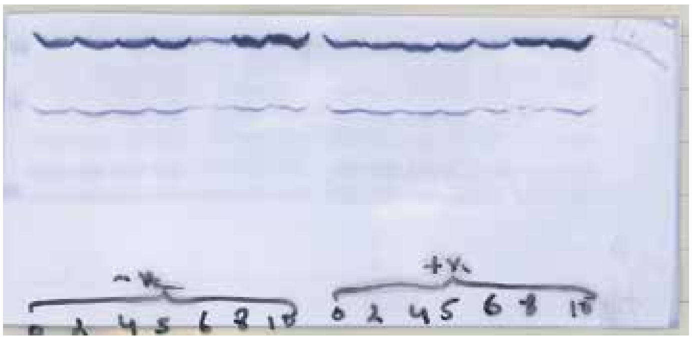

**Supplementary Figure S8.**

Uncropped images of western blotting in Figure 6a.
